# Data-independent acquisition mass spectrometry enables reproducible characterization of microbiota function

**DOI:** 10.1101/413021

**Authors:** Juhani Aakko, Sami Pietilä, Tomi Suomi, Mehrad Mahmoudian, Raine Toivonen, Petri Kouvonen, Anne Rokka, Arno Hänninen, Laura L Elo

**Affiliations:** Turku Centre for Biotechnology, University of Turku and Åbo Akademi, Turku, Finland; Turku Centre for Biotechnology and Department of Future Technologies, University of Turku and Åbo Akademi, Turku, Finland; Department of Medical Microbiology and Immunology, University of Turku, Finland; Department of Medical Microbiology and Immunology, University of Turku and TYKS Microbiology, Turku University Central Hospital, Turku, Finland

**Keywords:** Bioinformatics, Data-independent acquisition, Mass spectrometry, Metaproteomics, Microbiota

## Abstract

Metaproteomics is an emerging research area which aims to reveal the functionality of microbial communities – unlike the increasingly popular metagenomics providing insights only on the functional potential. So far, the common approach in metaproteomics has been data-dependent acquisition mass spectrometry (DDA). However, DDA is known to have limited reproducibility and dynamic range with samples of complex microbial composition. To overcome these limitations, we introduce here a novel approach utilizing data-independent acquisition (DIA) mass spectrometry, which has not been applied in metaproteomics of complex samples before. For robust analysis of the data, we introduce an open-source software package diatools, which is freely available at Docker Hub and runs on various operating systems. Our highly reproducible results on laboratory-assembled microbial mixtures and human fecal samples support the utility of our approach for functional characterization of complex microbiota. Hence, the approach is expected to dramatically improve our understanding on the role of microbiota in health and disease.

## Main text

Metaproteomics is an emerging research area, which aims to analyze an entire set of proteins from all microorganisms present in one ecosystem [1]. While the increasingly popular metagenomics provides insights on the functional potential of a complex microbial community, metaproteomics reveals the actual microbial function. In gut microbiota research, metaproteomics is expected to dramatically improve our understanding on the gut microbiota function and its role in health and disease [2].

The common approach in metaproteomics is to utilise data-dependent acquisition (DDA) mass spectrometry [3,4]. However, the performance of DDA has been reported to decline when sample complexity increases [5]. This may lead to limited reproducibility and dynamic range, bias toward high abundance peptides, and undersampling [5]. These are fundamental issues especially in comparative metaproteomics studies, in which reliable and reproducible quantification of peptides is crucial. It has been suggested that one possible approach to overcome these limitations is to use data-independent acquisition (DIA) modes of mass spectrometry [6], such as sequential window acquisition of all theoretical fragment-ion spectra (SWATH) [7]. The DIA techniques combine the advantage of the high throughput of shotgun proteomics with the benefit of the high reproducibility of targeted analysis, such as selective reaction monitoring [7,8]. However, DIA has not been applied to metaproteomics studies of complex samples before and currently there are no DIA data analysis software that could reliably process the data with the large sequence databases needed for metaproteomics.

To this end, we introduce here a DIA approach for metaproteomics together with a software package, named *diatools*, for analyzing the resulting DIA datasets. The data processing in the *diatools* package is divided into two major steps: 1) generation of the spectral library from DDA data, and 2) searching spectra from DIA data against the spectral library to identify and quantify peptides in the samples (Fig. 1). The output is a matrix containing the identified peptides as rows and their intensities for each sample as columns, which can then be further analyzed using various downstream data analysis tools. To ensure easy access to the software environment in a consistent and reproducible way, *diatools* was implemented as a Docker image, which is freely available at Docker Hub. Further details on the installation and usage of our *diatools* package are provided in the Methods section and in the software documentation (https://github.com/computationalbiomedicine/diatools).

**Figure 1.**
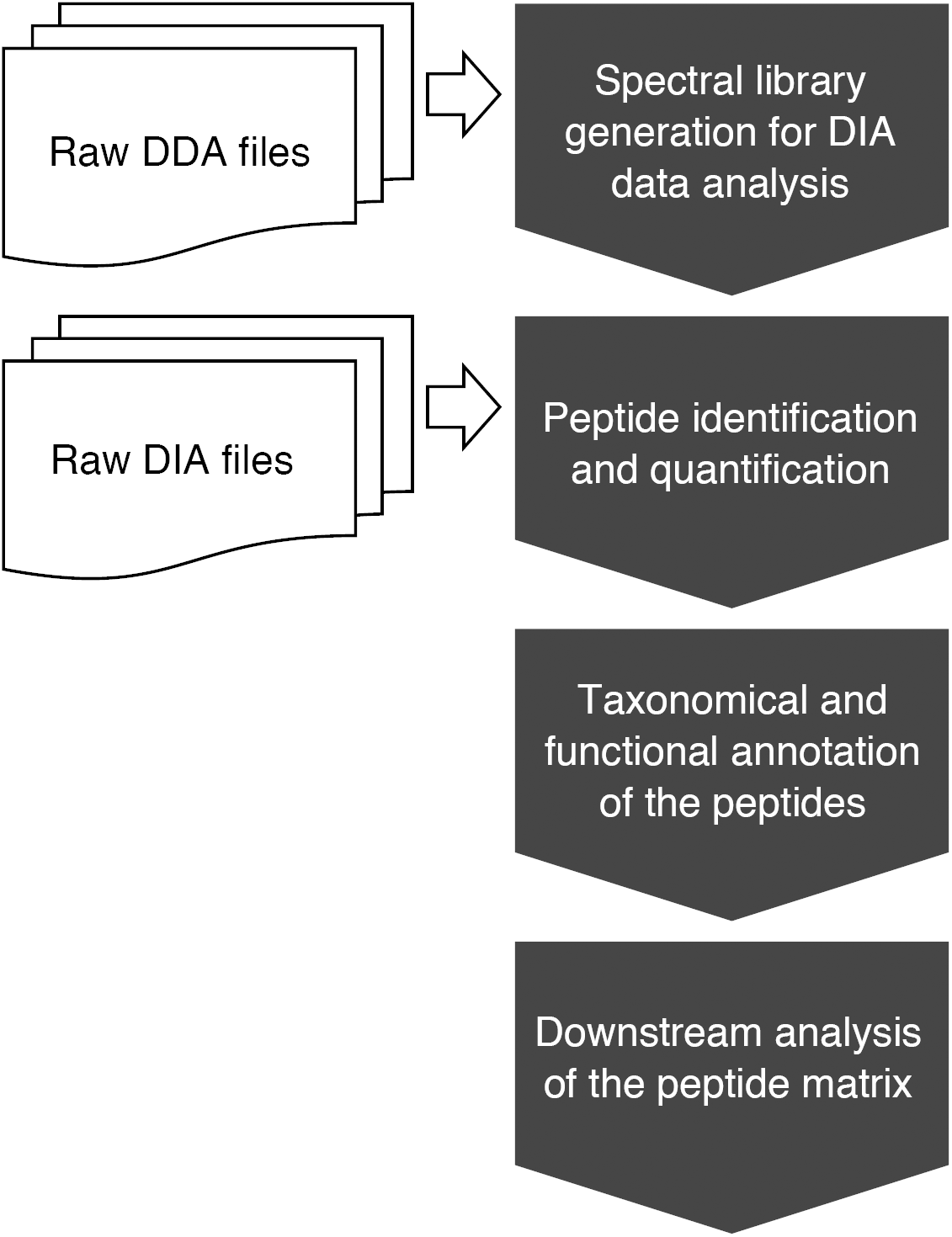
The workflow of our data-independent acquisition (DIA) approach for metaproteomics. The workflow consists of spectral library generation from data-dependent acquisition (DDA) data, and peptide identification and quantification from DIA data using the spectral library. After identification and quantification, the peptides are taxonomically and functionally annotated, and downstream analysis is performed on the peptide matrix. The data analysis workflow is implemented in the *diatools* software package.

**Figure 2.**
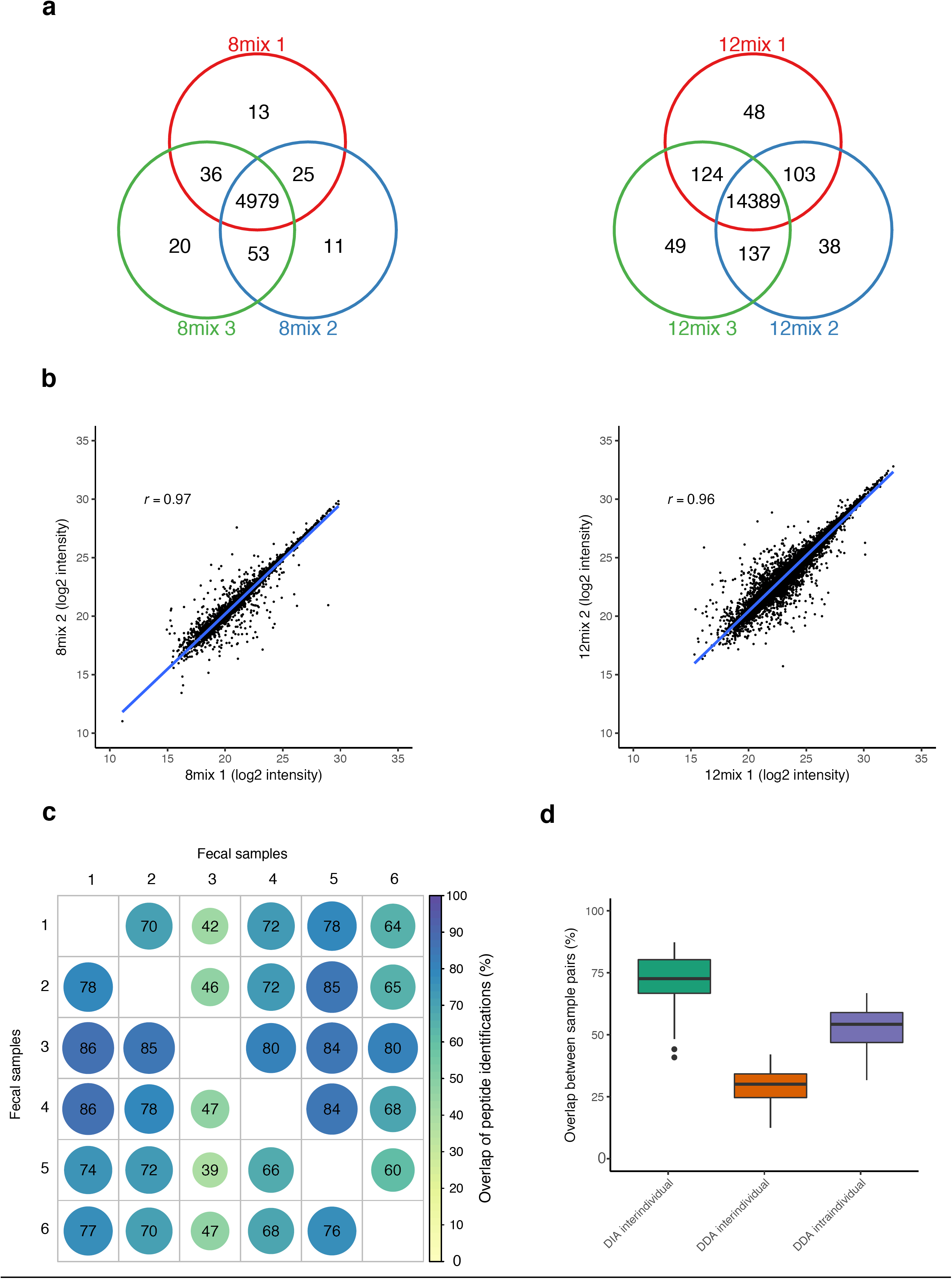
Reproducibility of data-independent acquisition mass spectrometry in metaproteomics. **(a)** Overlaps of the identified peptides between the three technical replicates of the 8mix (left) and the 12mix (right) samples. **(b)** Representative correlations of the peptide quantifications between two technical replicates of the 8mix (left) and the 12mix (right) samples; *r* = Pearson correlation coefficient. The rest of the pairwise correlations are shown in Supplementary Figure 1. **(c)** Overlaps of the identified peptides between each possible pair of human fecal samples. The proportions were calculated by dividing the number of common peptides by the total number of peptides in either of the samples of a sample pair, which results in an asymmetric matrix of percentages. **(d)** Overlaps of the identified peptides between all possible sample pairs within the current DIA study and a recent DDA study [10].

To evaluate the performance of our DIA metaproteomics approach, we first performed liquid chromatography-tandem mass spectrometry (LC-MS/MS) analysis of two mock microbial mixture samples containing eight and twelve different bacterial strains, referred to as 8mix and 12mix samples, respectively. Three technical replicates of both samples were analyzed. Additionally, we assessed six human fecal samples representing samples with a complex microbial composition. DDA data was used to generate the spectral libraries for the DIA data analysis, using the integrated reference catalog of the human gut microbiome [9] as the sequence database (detailed statistics of the spectral libraries are provided in Supplementary Table 1). The analysis of the DIA data identified then a total of 5137, 14888 and 12804 unique peptides in the 8mix, 12mix and human fecal samples, respectively (Supplementary Table 2).

Comparisons between the three technical replicates in the 8mix and 12mix data suggested that DIA had high reproducibility in terms of both identifications and quantifications when applied to metaproteomics samples (Fig. 1a-b). When comparing the overlaps of the identified peptides, more than 96% of the peptides were identified in all the technical replicates for both the 8mix and 12mix samples (Fig. 1a). Moreover, pairwise correlations between the intensities across the replicates were high (8mix Pearson correlation coefficient *r* = 0.97, 0.96 and 0.97; 12mix *r* = 0.96, 0.96 and 0.97) (Fig. 1b and Supplementary Fig. 1).

Also in the more complex human fecal samples, overlaps of the identified peptides across individuals were relatively high with the DIA approach; the median overlap of the identified peptides between sample pairs was 73% (interquartile range IQR 67% – 80%) (Fig. 1c-d). As a reference, in a recent DDA mass spectrometry gut metaproteome study of 16 healthy individuals over time by Kolmeder and co-workers [10], the intraindividual overlaps over time were 54% (IQR 47% – 59%) and interindividual overlaps only 30% (IQR 25% – 34%) (Fig. 1d). While the DIA approach is expected to reduce the problem of undersampling in metaproteomics [5], it should be noted that the pooled DDA samples used for building the spectral libraries may be affected by peptides that are commonly detected in the sample set. On the other hand, it should be also noted that the DIA data can be easily re-interrogated with updated or totally different spectral libraries, or the spectral libraries can be built for distinct purposes, such as to target specific proteins of interest or to fish out rare microbial proteomes.

To investigate the taxonomic characteristics of the peptides, we utilised the annotations of the integrated reference catalog of the human gut microbiome [9] and the lowest common ancestor algorithm (LCA) at phylum and genus levels. At phylum level, 76%, 71% and 47% of all the peptides detected were annotated in the 8mix, 12mix and human samples, respectively (Fig. 3a, Supplementary Table 2). At genus level, 51%, 43% and 28% of the identified peptides could be annotated in the 8mix, 12mix and human fecal samples, respectively (Fig. 3b, Supplementary Table 2). Importantly, only less than 0.1% of all the peptides were incorrectly annotated to phyla that were not in the 8mix and 12mix samples, and only 1% of the peptides were incorrectly annotated to genera not present in the 8mix or 12mix samples. Moreover, while the majority of the peptides of the human fecal samples remained unannotated, the taxonomic composition of the metaproteome was similar to those reported earlier by others [3,10,11].

**Figure 3.**
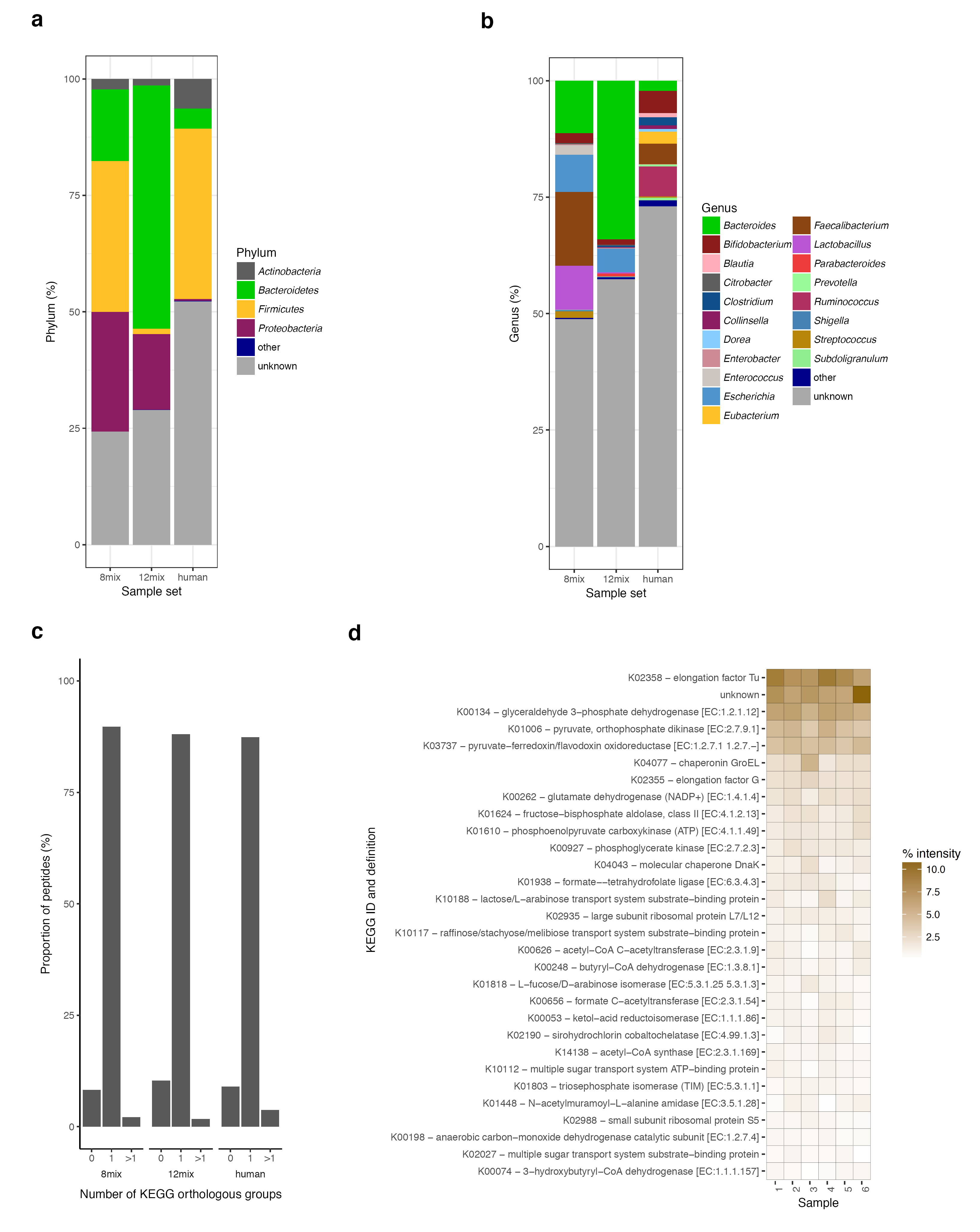
Taxonomic and functional characteristics of the peptides identified by the data-independent mass spectrometry workflow. Proportions of the identified peptides taxonomically annotated at **(a)** phylum or **(b)** genus levels. **(c)** Number of KEGG orthologous groups assigned to each peptide detected in the 8mix, 12mix and human fecal samples. **(d)** Heatmap of the thirty most common KEGG orthologous groups detected in the human fecal samples.

A major challenge in the taxonomic profiling of complex samples with both DDA and DIA metaproteomics is that different taxa may produce homologous proteins that share many identical tryptic peptides [6,12]. Therefore, LCA may introduce bias especially at lower taxonomic ranks. For instance, at genera level, taxa that have a large number of unique peptides tend to be overrepresented, while most peptides remain unannotated since they are shared between numerous genera [13]. Accordingly, our results indicate that when the microbial complexity of the sample increased, a lower proportion of the peptides could be annotated using LCA (Fig. 3a-b, Supplementary Table 2).

While the peptide-centric nature of bottom-up proteomics makes taxonomic profiling in metaproteomics challenging, the most important aspect of metaproteomics is to provide information on the functions of the microbiota. To investigate the functional characteristics of the peptides, we utilised the annotations of the integrated reference catalog of the human gut microbiome [9] and retrieved for each peptide all KEGG orthologous groups (KOGs) assigned to any of the proteins from which the peptide may have originated. Interestingly, a single KOG could be assigned to the majority of the peptides in all sample sets: 90%, 88% and 87% in the 8mix, 12mix and human fecal samples, respectively (Fig. 3c, Supplementary Table 2). This suggests that functional annotations can be assigned at peptide level without the intermediate step of protein inference and, hence, supports the use of peptide-based methods to detect functional differences in metaproteomics to avoid the potential ambiguities caused by protein inference. Similarly, peptide-centric methods have been previously suggested for single-organism proteomics [14].

In the human fecal samples of the present study, the most common KOGs included elongation factors, chaperones, and enzymes involved in glutamate production, glycolysis and butyrogenesis (Fig. 3d). These have been reported as typical microbial functions of the human gut microbiota by previous studies as well [10,11].

In summary, our results demonstrate for the first time that DIA can be applied to metaproteomics of complex samples and the results have good reproducibility. Similar to earlier studies that have used DIA for single-organism proteomics [7,8], a high overlap of identified peptides was observed across samples, enabling convenient downstream analysis of the data. Although the taxonomic profiling in metaproteomics remains challenging, our results suggested that the more important functional characterization of the metaproteomes was enabled by peptide-centric methods, which can avoid ambiguities related to protein inference. For robust analysis of the DIA metaproteomics data, we introduced the *diatools* software package, which is easy to use and customize for specific needs. The *diatools* package is the first open-source software specifically intended for DIA metaproteomics data.

## Methods

### The *diatools* software package

The *diatools* software package is designed for automatic analysis of data-independent acquisition (DIA) mass spectrometry data from raw data files to an intensity matrix that contains the identified peptides as rows and their intensities for each sample as columns. In short, d*iatools* first generates a spectral library from data-dependent acquisition (DDA) data and then uses it to identify and quantify peptides from the DIA data automatically. The first step was implemented according to the published protocol by Schubert et al. [15], while the second step builds on the OpenSWATH software [16]. The peptides are then taxonomically and functionally annotated and downstream analysis is performed on the peptide matrix.

For easy and reproducible access, the *diatools* package was implemented as a Docker image, which is freely available at Docker Hub repository compbiomed/diatools. Technically, the Docker image was built on the Ubuntu operating system (version 17.04) and contains the following preinstalled software: OpenMS [17] (version 2.3), Trans-Proteomic Pipeline (TPP) [18] (version 5.0), msproteomicstools [19] (version 0.6.0), ProteoWizard [20] (version 3.0.11252), and R [21] (version 3.3.2) with the SWATH2stats [22] (version 1.8.1). Step-by-step instructions to use the software are provided in the software documentation (https://github.com/computationalbiomedicine/diatools).

When developing the software package, we used a single server with 23 cores (Intel Haswell architecture). The available amount of RAM available in the server was 238 GB. In our datasets, the running time was 1 - 2 days per dataset. Additionally, we tested that all our datasets can be successfully run with 128GB of RAM.

### Samples

In the present study, two different laboratory-assembled microbial mixtures were used as sample material. The first of the two mixtures (8mix) contained an equal amount of eight different bacterial strains: *Bacteroides uniformis* ATCC 8492, *Bifidobacterium adolescentis* ATCC 15703, *Enterococcus faecalis* ATCC 700802, *Escherichia coli* K12, *Faecalibacterium prausnitzii* A2-165, *Lactobacillus acidophilus* ATCC 700396, *Staphylococcus aureus* NCTC 8325 and *Streptococcus pyogenes* serotype M6 ATCC BAA-946. The second microbial mixture (12mix) contained twelve different strains isolated from fecal samples of three human donors grown on fastidious anaerobe agar (LAB 090; LAB M, UK) and annotated by sequencing their 16S-rDNA: *Bacteroides vulgatus, Parabacteroides distasonis, Enterorhabdus* sp., *Bifidobacterium pseudocatenulatum, Eschericia coli, Streptococcus agalactiae, Bacteroides fragilis, Alistipes onderdonkii, Collinsella aerofaciens, Clostridium sordellii, Eubacterium tenue,* and *Bifidobacterium bifidum*.

Prior to mixing, the bacterial cell counts were equalized to 10 × 10^8^ cells / ml using flow cytometry (Bacteria counting kit for FLO, Fisher Scientific) and 1 × 10^8^ cells of each isolate were added to the final 8mix or 12mix, respectively. In addition, six human fecal samples from healthy anonymous individuals were analyzed under the permission of the Southwest Finland Hospital District.

### Protein isolation

The protein isolation for the 8mix and 12mix samples was performed using a Barocycler instrument NEP3229 (Pressure BioSciences Inc., South Easton, Easton, Massachusetts, USA), which uses pressure cycles to lyse the cells. From the human fecal samples, the proteins were isolated using the NoviPure Microbial Protein Kit (Qiagen, Hilden, Germany) following the manufacturer’s instructions.

Protein concentrations were determined using the Bradford method. Fifty μg of protein was used for trypsin digestion. The proteins were reduced with dithiothreitol (DTT) and alkylated with iodoacetamide. For the 8mix samples, pressure-assisted trypsin digestion was performed using the Barocycler instrument NEP3229 according to the manufacturer’s instructions. For the 12mix and human fecal samples, the trypsin digestion was performed conventionally in two steps: first trypsin was added in a 1:50 ratio and digested for 4h and then with a 1:30 ratio overnight at 37 °C. After digestion, the peptides were desalted using a SepPak C18 96-well plate (Waters Corporation, Milford, Massachusetts, USA).

### LC-MS/MS setup for data-dependent and data-independent analyses

The LC-ESI-MS/MS analyses were performed on a nanoflow HPLC system (Easy-nLC1200, Thermo Fisher Scientific, Waltham, Massachusetts, USA) coupled to a Q Exactive HF mass spectrometer (Thermo Fisher Scientific) equipped with a nano-electrospray ionization source. Five hundred ng of the digested protein samples were first loaded on a trapping column and subsequently separated inline on a 15 cm C18 column (75 μm × 15 cm, ReproSil-Pur 5 μm 200 Å C18-AQ, Dr. Maisch HPLC, Ammerbuch-Entringen, Germany). The mobile phase consisted of water with 0.1% formic acid (solvent A) or acetonitrile/water (80:20 volume/volume) with 0.1% formic acid (solvent B). A 90 min two-step gradient from 7% to 35% B, followed by wash with 100% B, was used to elute the peptides.

The MS data was acquired automatically using Thermo Xcalibur 3.1 software (Thermo Fisher Scientific). The DDA method consisted of an Orbitrap MS survey scan of mass range 375-1500 m/z followed by higher energy collisional dissociation (HCD) fragmentation for 15 most intense peptide ions. In the DIA method, the scan range was 400-1000 m/z.

For the DDA analysis, the peptide samples of each sample type were pooled and spiked with indexed retention time peptides (HRM Calibration kit, Biognosys, Schlieren, Switzerland). The pooled 8mix and 12mix samples were analyzed three times and the pooled human fecal samples six times in DDA mode. For the DIA analysis, each sample was injected once.

### Mass spectrometry data analysis

All data analysis steps from the raw DDA and DIA files to peptide intensity matrix were done using the *diatools* software package. The spectral libraries required for the DIA data analysis were generated separately for each sample type utilizing the data acquired in the DDA mode and two search algorithms: X!Tandem [23] (version 2016.01) and Comet [24] (version 2017.2.1.4). Parent ion mass tolerance was set to 10 ppm and fragment ion tolerance to 0.02 Da. The integrated gene catalogue [9] was utilized as the sequence database when performing the peptide to spectrum matching of the DDA data. The false discovery rate (FDR) was set at 1%.

The peptides were functionally and taxonomically annotated using the annotations of the integrated reference catalog of the human gut microbiome [9]. For each peptide, annotations of all possible target protein sequences were retrieved. A functional annotation for the peptide was assigned if the annotations of the target proteins were unambiguous. Similarly, a taxonomic annotation for a peptide was assigned at each taxonomic rank when possible using the lowest common ancestor (LCA) algorithm.

### Statistical analysis

The reproducibility of quantification between the technical replicates of the 8mix and 12mix samples was assessed by Pearson correlation.

## Abbreviations

DIA: data-independent acquisition
DDA: data-dependent acquisition
KOG: KEGG orthologous group
LC: liquid chromatography
LCA: lowest common ancestor algorithm
MS: mass spectrometry

## Declarations

### Ethics approval and consent to participate

The human fecal samples were collected from anonymous donors under the permission of the Southwest Finland Hospital District (reference number T254/2017).

### Consent for publication

Not applicable.

### Availability of data and materials

The mass spectrometry proteomics data have been deposited to the ProteomeXchange Consortium via the PRIDE partner repository with the dataset identifier PXD008738. The *diatools* software package source code is released under open source GPL 3.0 license and can be downloaded from GitHub https://github.com/computationalbiomedicine/diatools.git.

## Authors’ contributions

JA, SP, MM, PK, AR, and LLE designed the study. JA and SP wrote the first draft of the manuscript. RT and AH provided the clinical material and processed it together with PK and AR. JA, SP and TS wrote the software package. JA, SP, TS and MM performed the data analysis. LLE conceived and supervised the study and participated in writing the manuscript. All authors contributed to interpreting the results as well as editing the manuscript. All authors have read the final version of the manuscript and approved of its content.

## Acknowledgements

Not applicable.

